# Optimizing the multivariate temporal response function(mTRF) framework for better identification of neural responses to partially dependent speech variables

**DOI:** 10.64898/2026.02.25.707435

**Authors:** Konrad Dapper, Sarah Hollywood, Taylor Dool, Blake Butler, Marc Joanisse

## Abstract

An increasingly popular approach to investigating the neural bases of speech processing is forward modeling via a multivariate temporal-response function (mTRF). This approach uses stimulus characteristics to predict neural responses, especially in EEG and MEG. A central question in this regard is how best to represent the input stimulus. In the case of speech processing, established representations include the speech envelope or spectrogram, as well as feature-based linguistic representations of phonetic content. However, when multiple representations are used as input, a key challenge is how best to isolate their relative effects. This is particularly challenging because such representations have nonvanishing mutual information. To address this problem, we propose optimizations to the mTRF framework via a novel statistical approach of cyclic permutation. Additionally, we propose methodological improvements to the mTRF model targeting three key challenges: effectively managing spatial and temporal autocorrelations endemic to multi-sensor EEG data; mitigating the effects of endogenous drift; and introducing robust artifact rejection to enhance data quality. To demonstrate the effectiveness of this approach, the novel method was applied to a novel EEG data set of natural language listening in 27 adults with normal hearing. Our data showed that including ICA decomposition, artifact rejection, and cyclic permutations in an mTRF analysis improves the isolation of neural responses specific to phonetic and acoustic input variables.

**Author Summary:** Speech processing happens in different stages. It starts with recognizing basic sounds, then categorizes them into discrete categories called phonemes, and goes on to understanding words and sentences. The multivariate temporal response function (mTRF) is a method for predicting brain activity from different features of the speech stimulus. Features that can be used as input to the mTRF model include acoustic features, such as sound envelopes, as well as more abstract language features, such as phonemes, which are a fundamental building block of words. One problem in speech research is distinguishing neural responses to different features. This is challenging because knowing one feature of the speech stimulus enables educated guesses about others and educated predictions about how this feature will behave in the future. Both of these properties of speech make multivariate temporal statistical analysis more difficult. To address this, we propose changes to the preprocessing of the EEG recordings and a new mathematical model that uses a partially rearranged version of the features of the speech stimulus to isolate the predictive power of a particular type of speech feature.

## Introduction

Speech perception is a vital part of human communication and can be investigated using Electroencephalography (EEG) or magnetoencephalography (MEG). Using these and other techniques, deficits in speech processing have been shown to contribute to a wide range of neurological pathologies, including developmental learning disorders [1–3], autism [4,5], and presbyacusis [6,7].

Typically, EEG and MEG tasks investigating speech processing have relied on event-related designs involving a series of repeated individual trials and exploring the neural responses to those trials. A benefit of this approach is that it enables significant control over the stimulus, but a downside is that the paradigm becomes repetitive and induces relatively non-contextual processing of individual trials, as repeated trials differ from how speech is encountered day-to-day.

An emerging alternative method is to investigate responses to speech via models that continuously predict neuroelectric activity from ongoing speech stimuli. This approach benefits from the option to use natural stimuli such as spoken narratives, newscasts, and stories, which are generally more engaging and naturalistic. But unlike trial-based designs, where averaging responses across multiple trials is a straightforward way to improve the signal-to-noise ratio, processing responses to natural speech requires more complex modeling to isolate effects of individual linguistic elements of interest (e.g., phonemes, morphemes, words, phrases). In the literature, these models are collectively referred to as forward models [8–11].

When choosing the input to such a model, a natural first approach is to use a time trace of speech derived from its acoustic profile. And indeed, the literature clearly supports the view that the primary factors driving the neural response to speech are the amplitude envelope and the spectrogram of the input [12–14]. After all, similar evoked responses to those observed during speech processing can be seen even for stimuli that, despite being acoustically similar to natural speech, do not convey any meaning, such as backward speech [12] or responses evoked by an actively ignored speaker [13]. However, a model based solely on acoustic input variables is incomplete, as it lacks any description of how the signal is broken down into its constituent linguistic units and its meaning is subsequently comprehended. A complete model of speech processing must include these aspects. This can be seen by, for example, specific neural responses to speech processing, as shown with auditory chimes [14], or differences in responses to speech and music [15].

Modeling of speech in a way that captures the full breadth of its features is a complex task. Speech features span multiple time scales, from the identification of the fundamental frequency of a speaker at ∼100 Hz [16], to slower processes like phoneme identification (duration ∼100 ms), or the understanding of words and sentences, which can take on the order of a second or even longer. And indeed, many of these different hierarchical layers are predictive of the neural electrical responses evoked by speech [15,17,18].

The mTRF framework [8] approaches this problem by allowing for different types of input mappings either concurrently or within separate models. The central assumption is that the weighted sum of stimuli within a time window around the observation time point can predict subsequent neural activity, with each weight acting as a parameter to be determined via fitting of a linear model. A common issue with high-dimensional models of this sort is overfitting, given the explosion in parameters inherent to mapping physically complex stimuli like speech to multi-channel, time-varying neural signals like EEG. To minimize this issue, the mTRF approach employs ridge regression [8], which reduces overfitting by assuming all parameters are likely to be small, with a ridge parameter λ controlling the extent of this assumption.

One of the key benefits of mTRF is versatility. The same modeling framework can be used to investigate multiple aspects of speech perception and comprehension processes by choosing different input variables, such as the spectrogram, phonetic features, word onsets, or linguistic surprisal. However, a fundamental challenge is that many of the input spaces (e.g., the phonetic features of speech and its spectral characteristics) are not statistically independent. Or, in simple terms, knowing the phonetic features of speech allows one to predict a significant amount of the auditory spectrogram. This conflict is typically addressed by comparing models that differ in the input variables they include [19,20]. While this is a valid approach, we demonstrate here that it can be improved by better controlling for overfitting, which remains an issue when comparing models with different input sizes despite the use of ridge regression. We propose that this additional uncertainty can be managed by introducing cyclic permutation statistics.

A prerequisite to this novel statistical approach is the implementation of three useful optimizations to the mTRF framework. As we demonstrate, this combination of refinements enables improved identification of neural activity evoked by individual aspects of speech processing and addresses three potential weaknesses. The first is that the regularization of the mTRF via ridge regression implicitly assumes that the EEG channels are statistically independent. However, in general, neighboring EEG channels are correlated, violating this assumption. This problem is rectified by performing the modeling using ICA components rather than the EEG channels as inputs, since ICA components are stochastically independent by construction. The second issue is that most mTRF implementations we are aware of [8] leave aside the stabilization of endogenous drift and artifact rejection. Here, endogenous drift refers to all possible slow changes in the participant’s recorded neural state that the input variables cannot capture, such as vigilance. Both endogenous drift and artifact rejection are addressed by altering how the data is partitioned during model training. Third, the ridge parameter λ is generally determined with a k-fold cross-validation procedure, which is relatively computationally intensive and, as we demonstrate below, susceptible to noise. Therefore, we have replaced it with a numerical simulation of the ridge parameter, with the benefit of greatly improved computational efficiency.

In this study, a total of 27 participants were recruited (12 male, 14 female, and one person self-identifying as non-binary). Participants performed an active listening task while their neural activity was recorded using electroencephalography (EEG). During the experiment, participants listened to six audio stories, each approximately 6 minutes long, followed immediately by six comprehension questions per story to ensure vigilance. In total, three participants were excluded: two for not completing the EEG measurements and one for poor EEG signal quality (more than 50% of stories rejected during the analysis), resulting in a final cohort of 24 participants (10/13/1 m/f/nb). The mean age of the included participants was 23.1 ± 5.9 years (min = 18, max = 40). All had normal hearing, verified by pure-tone audiogram measurements at 0.5, 1, 2, and 4 kHz with a maximum threshold of 20 dB NHL for both ears independently.

## Results

### EEG results

#### Evaluating the Relative Importance of Spectral and Phonetic Features Using the Conventional mTRF Model

We first computed a conventional mTRF model of the dataset. That analysis was performed in EEG channel space, with an optimal lambda calculated for each participant using five-fold cross-validation, and data segmented into six segments, each ∼60 seconds in duration (five for the cross-validation to determine the ridge parameter and one for the final validation). This analysis was conducted independently for each participant and story.

Figure 1 shows both the average of all correlations (gray bars, panels A-C) and the mean correlation observed for each individual story (colored bars/lines). The first column of Figure 1 (panels A, D) shows the correlation coefficient when only the spectrograms are used as input to the mTRF model. In comparison, the middle column (panels B, E) shows the results when only the phonetic feature vectors are used as input to the mTRF, and the third and final column (panels C, F) shows the result of a model using both as input.

**Figure 1.**
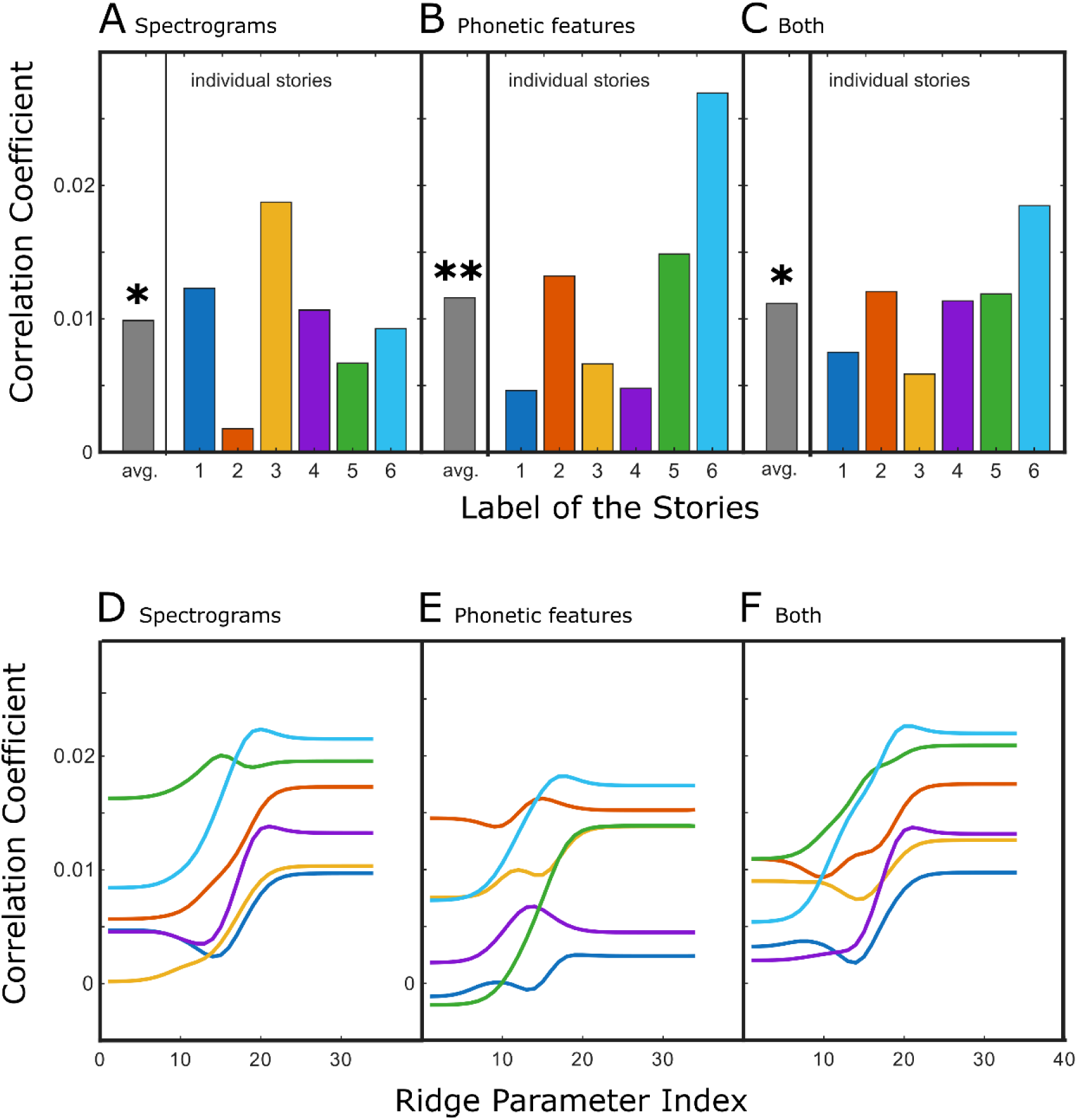
**Correlations analyzed using the conventional mTRF model**: **In the top row**, bar charts showing the mean correlation coefficient of the best ridge parameter. **In the bottom row**: the correlation coefficient as a function of the ridge parameter index. A, D, using only the spectrogram as input to the mTRF model. B, E: only phonetic features as input; C, F: both as input. The colors in the charts represent the different story conditions: gray indicates the combined data from all stories, dark blue represents the ’funny’ story, orange corresponds to ’fantasy,’ yellow to ’boring,’ purple to ’realistic,’ green to ’adventure,’ and light blue to ’boring’ story 2.

To statistically evaluate the models, we first tested whether the correlation coefficients are different from zero using a two-sided t-test against a constant model. We found that in all conditions (i.e., using only spectrograms, only phonetic features, or a combination of the two), there was a significant effect (t(135)=2.2/2.8/2.6); p = 0.0294/0.0055/0.0103for the average across all stories (Figure 1A, B, and C). Paired t-tests further showed no difference among model correlation coefficients (A vs. B: t(135) = 0.612, p = 0.7299; A vs. C: t(135) = 0.218, p = 0.5858; B vs. C: t(135) = 1.378, p = 0.9146).

The statistical analysis described above was repeated for each story independently, without finding any statistically significant effect, either for a model against zero or for comparisons between models, indicating that the SNR of a single story is insufficient.

One factor that might have limited the model’s performance is the substantial variation in the optimal ridge parameter (λ) across stories and its impact on the correlation coefficient. To visualize this problem, Figure 1 D-F shows the mean correlation coefficient as a function of the ridge parameter. Our data show substantial variation in the distribution and optimal λ values between the different stories.

#### Does the introduction of ICA and finer segmentation improve the reliability of the mTRF model?

The conventional mTRF model explored above exhibits uncertainty in selecting the optimal ridge parameter (Figure 2 A,D). When investigating the effects of the transition into ICA space and artifact rejection on the lambda parameter, our data show that the maximal slope of the correlation coefficient as a function of lambda differs across the different models. Both the transition into ICA space and the introduction of artifact rejection decrease the spread of the optimal lambda. This was statistically verified using a Kolmogorov-Smirnov test (Figure 2 D vs. E: ks2 = 0.30, p = 5.4233e-06; Figure 2 E vs F: ks2 = 0.20, p=0.0044).

**Figure 2:**
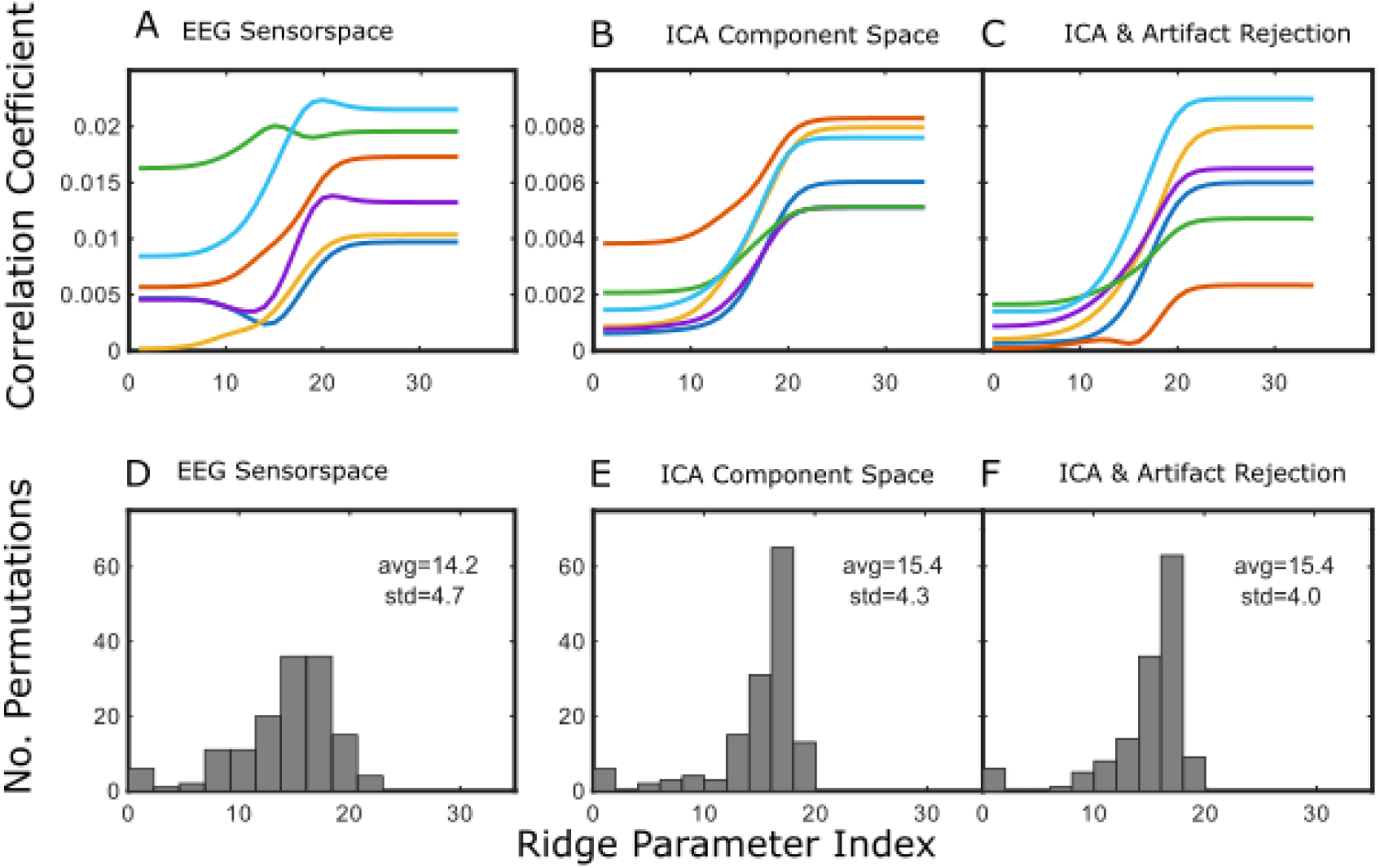
distribution of the ridge parameter lambda: The figure illustrates a comparison of the reliability of cross-validation in determining optimal lambda values across three methodological approaches. Subplots A and D display results using the conventional mTRF algorithm; B and E present the conventional mTRF in ICA-transformed space; and C and F showcase the conventional mTRF algorithm in ICA space with small data partitions and automated artifact rejection. The top row presents the mean correlation coefficient plotted against the ridge parameter (lambda), providing a clear depiction of how model performance varies with regularization strength. Each color in the plot represents a specific story. Their colors follow the same convention as in Figure 1

Another point to note is that, at the given noise level, the correlation coefficient remains relatively flat for higher lambda values, which aligns with our simulation results. As a result, cross-validation performed for individual participants is quite susceptible to noise in specific recordings—a problem that can be mitigated by simulating lambda.

#### Evaluating the Relative Importance of Spectral and Phonetic Features Using the Optimized mTRF Model

We next evaluated the performance of the optimized mTRF model (Figure 3). The first row of the figure shows a histogram of all cyclic permutations, with the red bracket indicating the part of the correlation that can be attributed to a given set of input variables. The second row shows the correlation that can be attributed to the given input variables, averaged across stories (gray) and for each story individually (colored bars).

**Figure 3:**
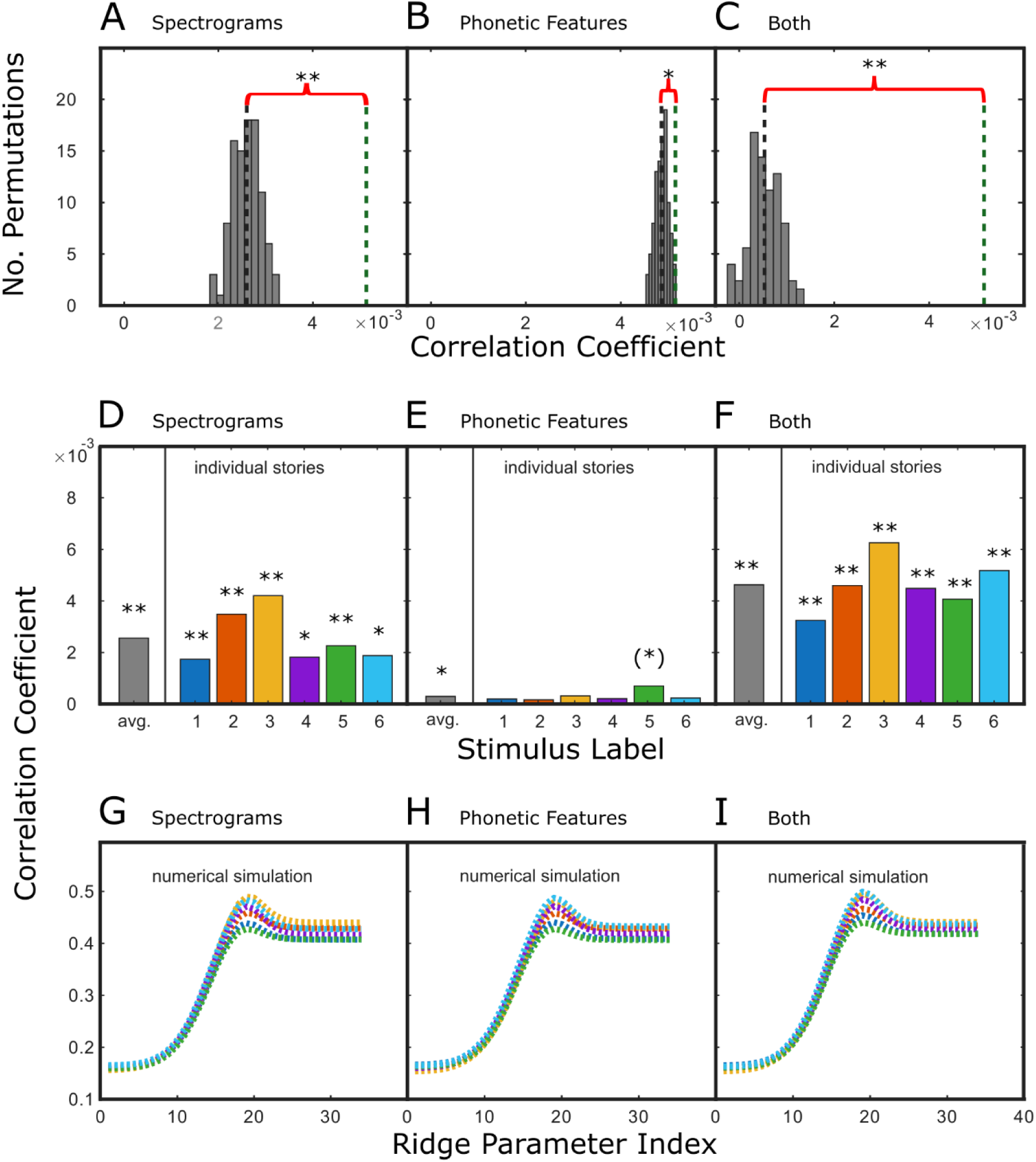
Correlations analyzed using the optimized mTRF model: This figure presents the correlation coefficients for Spectrograms (A, D, G) and phonetic feature vectors (B, E, H), as well as the sum of both (C, F, I). **The top row** (A–C) displays histograms of the average correlation coefficient for all stories across all permutations. The red bracket visualizes the correlation coefficient assigned by the model to the specific component. **The middle row** (D–F) shows the mean correlation coefficients. The first bar (gray) represents the overall mean across all stories, while the subsequent bars represent individual stories: Each color in the plot represents a specific story, following the same convention as in Figure 1. For each story, N = 136 (24/24/22/24/20/22), based on the rejection criterion described in the Methods section. Note. *p(corr) < 0.05, **p(corr) < 0.01, (*)p<0.05 (uncorrected). **In the bottom** row, the simulated mean correlation coefficients for the different stories and lambda values (same color convention as in the middle row)

Figure 3 A shows the difference between the correlation explained by all variables and the correlation that remains if the spectral information is removed from the input space via cyclic permutation (p<0.001). Figure 3 B shows the correlation that can be attributed to the phonetic feature vectors. The results demonstrated that the addition of phonetic feature vectors improves the auditory model (p = 0.018). Our data show that even when all stimulus channels are cyclically permuted (Figure 3C), a non-zero correlation remains, demonstrating some model overfitting, which the novel model compensates for.

Beyond the global picture on the importance of the auditory spectrogram for mTRF modeling, we also examined responses to individual stories. Our data revealed that the spectrograms could enhance the explained correlation beyond what the phonetic features could explain, with p-values of 0.003, < 0.001, < 0.001, 0.014, 0.003, and 0.015, respectively, for the six different stories. As such, across multiple stimuli, the inclusion of the spectral information could explain more correlation than the phonetic features alone. In contrast, the addition of phonetic features generally failed to improve the model’s performance on individual stories, with only one story reaching statistical significance (p = 0.025), which did not survive false discovery rate correction (p < 0.008 given 6 comparisons). The model in which all channels were cyclically permuted resulted in p-values for all individual stories less than 0.001.

In summary, our model showed that combining phonetic features and spectrograms is more effective for predicting the neuroelectric response to speech. Additionally, the spectrogram proves to be more critical for the model than the phonetic feature vector. Indeed, the fact that this effect persisted in individual stories indicates that shorter recording periods provided sufficient SNR to isolate the impact of individual components of the input to the model.

#### Comparing the conventional and the optimized mTRF model

One way to compare the conventional and the optimized mTRF models is to analyze the unique and combined explanatory contributions of each set of input variables. This approach enables us to examine the specificity of the models. Here, we compute these contributions by dividing the median ratio of the sum of the correlations explained by the two smaller models by the correlation explained by the model using both inputs simultaneously. Using the conventional mTRF model, we find a median ratio of 1.27 (Figure 4A), indicating that the sum of the correlation coefficients from the two separate old models—one using only the spectrogram and the other using only the phonetic feature vectors—explains 27% more correlation than a model using both inputs together. In contrast, the same median ratio computed using the new model was 0.77, indicating that the two channels, when considered separately, accounted for only 77% of the correlation.

**Figure 4:**
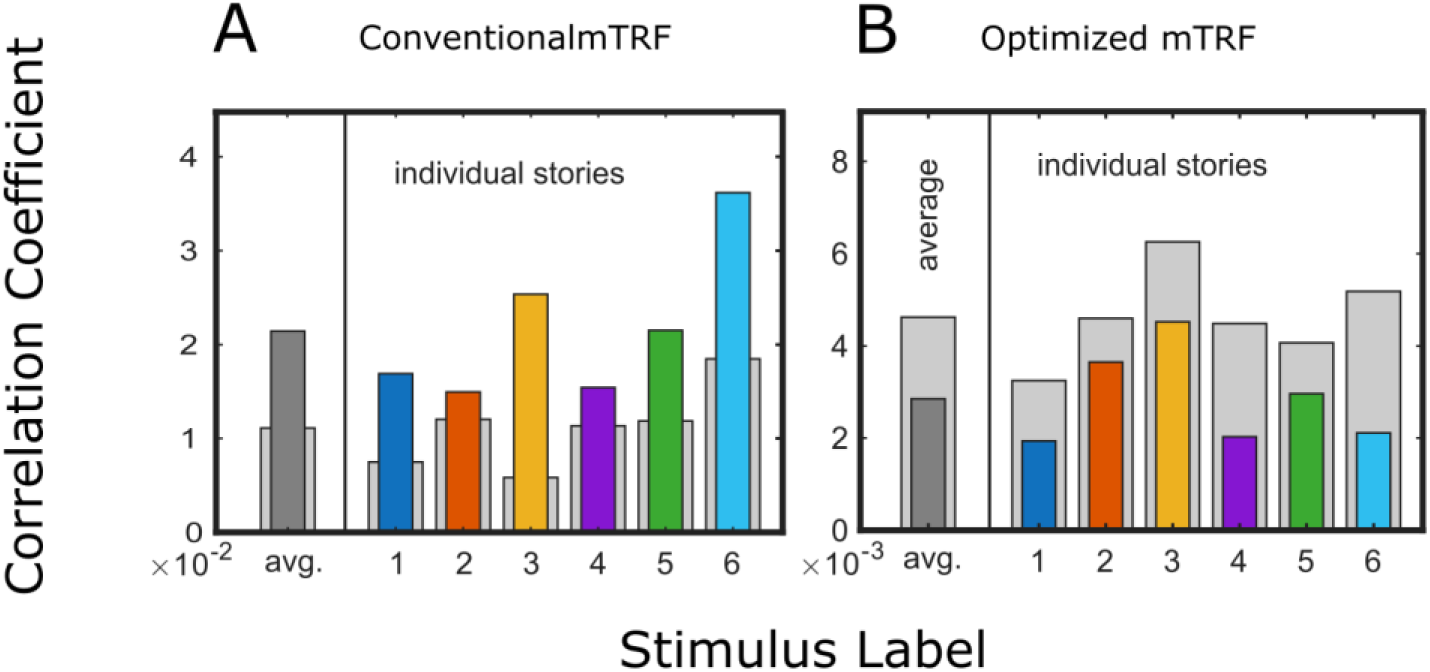
Comparison of conventional and optimized MTRF. **This** figure compares the effects of different sets of input variables. Colored bars represent the sum of correlations explained by either the spectrogram or phonetic variables as independent inputs, while the broader light gray bars show the total correlation coefficient achieved when both the spectrogram and phonetic variables are used simultaneously. The colors correspond to the different stories, following the same convention as in previous plots. Panel A presents results from the conventional mTRF model, while Panel B presents results from the optimized mTRF model.

This comparison demonstrates that the conventional model is constrained by mutual information, as a significant portion of neural responses cannot be attributed exclusively to either phonetic or auditory features. In contrast, the optimized model eliminates this overlap between the two sets of variables. These observations are statistically significant, as the ratios were drawn from different distributions, as determined by a Kolmogorov-Smirnov Test, with p-values < 0.001 for all comparisons.

## Discussion

This study used EEG recordings from 27 normal-hearing adults, obtained during active listening to natural speech (six short stories), to investigate optimizations to the mTRF model. To optimize the mTRF model, we performed ICA-based analysis, fine-grained data partitioning, and introduced a novel statistical method (cyclic permutations) to isolate neural activity exclusively evoked by speech’s phonetic or auditory features(spectrograms). We focused on phonetic features and spectrograms for the evaluation of the novel approach, as the literature suggests they independently contribute to speech processing [3,19], while also sharing a significant amount of mutual information; this makes them ideal candidates to investigate the performance of the optimized mTRF model.

There were two key findings. First, the sensitivity of the mTRF model was improved by the three alterations to the conventional approach. Second, these optimizations also improved the model’s specificity in attributing independent variance to the input variables.

### Sensitivity of the conventional and optimized mTRF model

The optimized mTRF model was more sensitive than the conventional mTRF model. Consistent with the literature, we observed that both spectrograms and phonetic features—analyzed separately or in combination—accounted for a significant proportion of the EEG responses in our participant cohort when analyzed using a conventional mTRF model. However, these general findings could not be confirmed when examining responses to individual stories (Figure 1). In contrast, the optimized mTRF model not only confirmed these findings, but also identified a significant contribution of the spectrogram or both input variables to the correlation in responses to individual stories (Figure 3).

The conclusion that the optimized mTRF model is more sensitive than the conventional mTRF model may appear to be undermined by the observation that the total correlation strength is weaker in the optimized model (Figure 3) than in the conventional model (Figure 1). However, two factors can explain this apparent discrepancy.

The first is that at least part of the explained correlation observed in the traditional model is due to overfitting. In contrast, the optimized mTRF model compensates for model overfitting and correlations that are not specific to a particular set of input variables by subtracting the surrogate data’s mean correlation. This necessarily decreases the correlation strength.

The second argument is that the optimized approach is conservative, in that the reported findings are averages across all ICA components. However, it’s highly improbable that all ICA components are actually related to speech processing; consequently, averaging dilutes the signal, making an underreporting of the total correlation strength very likely. One might argue that a similar dilution effect occurs in sensor space, due to the inclusion of electrodes that likely capture little speech-evoked neural signal; however, given that speech processing across different regions is expected to be statistically dependent, ICA is likely to compress the neural responses evoked by speech into only a handful of statistically independent components. Thereby, the normalization of all ICA components likely amplifies the dilution effect in ICA space. The approach of averaging all ICA components was chosen to ensure that no part of the neural response is lost during the analysis. Future studies may refine this approach by selecting specific subsets of the ICA components.

At a conceptual level, two factors explain why our proposed optimization to the mTRF model improved sensitivity. The first is reduced noise susceptibility through artifact removal and stabilization of endogenous drift; this is essential, as factors such as participant movement and inattention alter the neural response to speech stimuli in ways that are difficult to predict [13,21,22]. Here, we minimize the effect of these variables on the observed correlation by ensuring that training and validation samples are evenly distributed in time. This ensured that both the training and testing of the model uses data with an average level of attention, motion artifacts, and other endogenous drift factors that affect speech processing or EEG measurements.

The second factor contributing to improved sensitivity is a more stable estimation of the ridge parameter. All stories in our stimulus set had very similar acoustic and phonetic properties, and thus participants’ neural responses would also be expected to be similar. Numerical simulations showed that, in these conditions, the optimal ridge parameter for the different stories is expected to have a limited spread. And yet, we observed considerable variation in the ridge parameter determined by the conventional mTRF model (Figure 2). This suggests that noise is interfering with the k-fold cross-validation process used to determine the optimal lambda. We found that transitioning from EEG channels to ICA components decreases the spread of the ridge parameter (Figure 2). This is likely due to the fact that the ICA-based data better meet the statistical independence of observations that is a prerequisite of ridge regression [29]. Furthermore, the spread of the ridge parameter is reduced by artifact rejection, likely because it improves overall data quality. In summary, these technical improvements and the transition to simulated ridge parameters decreased model uncertainty by reducing the uncertainty in the ridge parameter, thereby improving the model’s sensitivity.

### Specificity of the conventional and optimized mTRF model

A second key benefit of the optimized mTRF model is that it increases specificity. The compensation for the surrogate data set’s mean correlation ensures that the optimized mTRF model can directly measure correlation specific to a subset of the input variables. Consequently, the observed coefficients shown in Figure 3 A, B are direct measures of the contribution of individual speech features. In contrast, when trying to measure these effects using the conventional model, we have to rely on the indirect approach of comparing the correlations of different models using t-tests. In the current study, t-tests could not resolve these differences between models.

At the conceptual level, three factors contribute to this increased specificity of the optimized mTRF model. The first is that, for the reasons outlined above, the optimized model has increased sensitivity that helps resolve minor differences across different input spaces. The second factor is that the conventional mTRF is expected to exhibit greater overfitting-induced variability when comparing different input features than the optimized model. In the conventional approach, when comparing two features that are subsets of the space of input variables, two different models must be fitted. A downside of this approach is that fitting two models with varying input space size increases overfitting-induced variability. In contrast, when using cyclic permutation, the same total input space is always present; thus, overfitting potential is equal in all conditions and does not impede the analysis. Finally, the third factor is that the auditory and phonetic input variables have some degree of mutual information. In the conventional mTRF model, this duplication hinders clear interpretation since comparing individual model fits to the combined models fails to identify the unique contribution of one or the other. The presence of the overlap can be seen by the fact that the sum of the two traditional models in the present study explained a median of more than 100% (127%) of the total explained correlation. In contrast, the optimized mTRF model eliminated the variability introduced by mutual information by only measuring the unique variability of each subset, as can be seen by the fact that the sum of the two subspaces accounts for a median of less than 100% (77%) of the total correlations explained by the sum of both individual correlations (Figure 4).

### Limitations of the mTRF model

Any model is only applicable in specific circumstances; therefore, it is essential to acknowledge the limitations of mTRF models and of the novel statistical approach. First, all mTRF models are linear and thus cannot capture nonlinear interactions among input variables. This restriction may limit the model’s ability to reflect the full complexity of neural processing. However, this limitation can be at least partially overcome by identifying the input variables that best linearize speech processing. Another limitation is that mTRF models do not allow inferences about non-phase-locked neural activity, or, in other words, activity induced by the stimulus. Together, these could help explain why all mTRF models typically account for only about 1% of neural activity. In addition to these general limitations of the mTRF model, one should keep in mind that the cyclic permutation analysis captures only additional evoked neural responses. Therefore, a lack of correlation between the mTRF model and the input subset does not imply that a model using only this subset of input variables cannot predict neural responses to speech.

Another limitation is the spacing of the cyclic permutations, which here is approximately 4 seconds. The minimum spacing is a function of the length of the stimulus and the desired number of permutations. While a spacing of 4 seconds is sufficient for investigating responses to acoustic and phonetic features, future studies investigating lexical, semantic, and contextual effects (i.e., speech features with longer autocorrelation lengths) may require longer stories to ensure that the time spacing of the cyclic permutations is longer than the autocorrelation of the speech features analyzed.

Finally, we note that the optimizations proposed here should generalize well to other approaches to linking neuroelectric data to speech. For example, an alternative approach to regularization has been proposed in which the mTRF kernel is determined via the Boost algorithm [29] - a variant of gradient descent. There is no conceptual reason to think that analysis in ICA space, smaller partitioning, and cyclic permutations could not be adapted to this algorithm. We do note, however, that the analysis using the Boost algorithm relies on a predefined hierarchical order of features [29], which is not required by the approach presented here.

Looking ahead, future research can use these methodological improvements to address several aspects of speech processing. One option is to replicate these measurements in cohorts expected to exhibit reduced phonological processing efficiency. Potential groups include non-native speakers of a language, individuals with cortical hidden hearing loss [6], the elderly [23], children [24], and those with developmental language disorders [25]. This is a promising avenue, as all these populations could benefit from improved model specificity in identifying neural activity associated with the type of phonetic classification presented here. Likewise, the novel statistical approach to the mTRF framework is not limited to questions concerning phonological features, but could also be generalized to investigate neural responses associated with higher-level linguistic and contextual cues or even multisensory inputs.

## Methods

### The conventional mTRF model

The conventional mTRF model used the mTRF-Toolbox[8] (https://github.com/mickcrosse/mTRF-Toolbox.git). Input variables were used to predict the EEG signal at a 64Hz sample rate, with a window that started 600 ms before stimulus onset and ended 200 ms after the observation point, for a total of 53 time points. A ridge regression model was trained with k-fold cross-validation on k-1 folds of the data and applied to a series of trial parameters to determine the optimal value of the ridge parameter (λ). Specifically, we used k = 6, and 34 lambda values, spaced logarithmically from 10^ (−2) to 10^9.

### Optimized mTRF model

The proposed statistical framework modifies the conventional mTRF code and is available for download under a Creative Commons license (https://github.com/ugedj/cyclic-mTRF). It departed from conventional mTRF models in several aspects.:

**1) data partitioning:** The conventional mTRF model involves segmenting the data into ∼6 segments for cross-validation to determine the optimal lambda (which, in this study, results in 60-second-long epochs). It does not contain artifact removal[8], likely because artifact removal introduces gaps in the data that complicate the implementation of mTRF analysis. We propose optimizing this by splitting the data into many shorter epochs (1 second long in this study). Using shorter epochs allows artifacts to be identified and removed based on the signal’s variance. The remaining segments are then assigned sequentially to the k different folds. At the same time, ensuring that each fold contains the same number of epochs. A formal explanation of why this is possible is provided in the supplementary information.

A downside of this approach is that, because both the EEG and the speech signal are autocorrelated, short epochs introduce an additional risk of overfitting in the model, which must be accounted for in the statistical analysis of the results.

**2. Determining the optimal ridge parameter via numerical simulation.** As noted above, cross-validation is typically used to determine the optimal ridge parameter (λ). Using numerical simulation instead comes with the benefit of a reduction in the noise susceptibility of the lambda calculation. The algorithm also reduces computational effort by 95%, making statistical modeling feasible on ordinary computer hardware.
**3. Fitting the model in ICA space.** Linear models are typically fitted in EEG channel space. As noted above, the spatial correlation of EEG data, especially in dense channel arrays, introduces correlations between neighboring channels that violate key assumptions of mTRF models. Computing the mTRF in ICA space ensures statistical independence among EEG channels, which is more appropriate for the ridge regression method, as it is a prerequisite for simulating lambda.

### Addressing effects of EEG autocorrelation as a source of overfitting

Permutation analysis is a widely used method for evaluating the statistical significance of neural electrical measurements. In essence, this approach involves randomly reassigning stimuli to recorded epochs to generate surrogate-datasets. These surrogate-datasets serve as a reference for estimating the likelihood that the observed results could have occurred by chance.

The permutation analysis approach makes a key, yet often unspoken, assumption that shuffling the stimuli does not alter the fundamental statistical properties of the experiment. In simple terms, this means that if the measurement were repeated with a shuffled stimulus, participants would show identical neural and behavioral responses. However, this assumption is violated when using natural speech, as shuffling disrupts the inherent temporal structure and autocorrelation present in natural speech. To address this, the permutation algorithm must be modified so that the surrogate-stimuli retain the same autocorrelation properties as the real stimuli. One effective solution is to use cyclic permutations of the stimulus (see Figure 5). Here, a cyclic permutation means the analysis assumes the stimulus begins at a point other than the actual start of the sound. When the sound reaches the end, it simply cycles back to the beginning of the stimulus.

**Figure 5:**
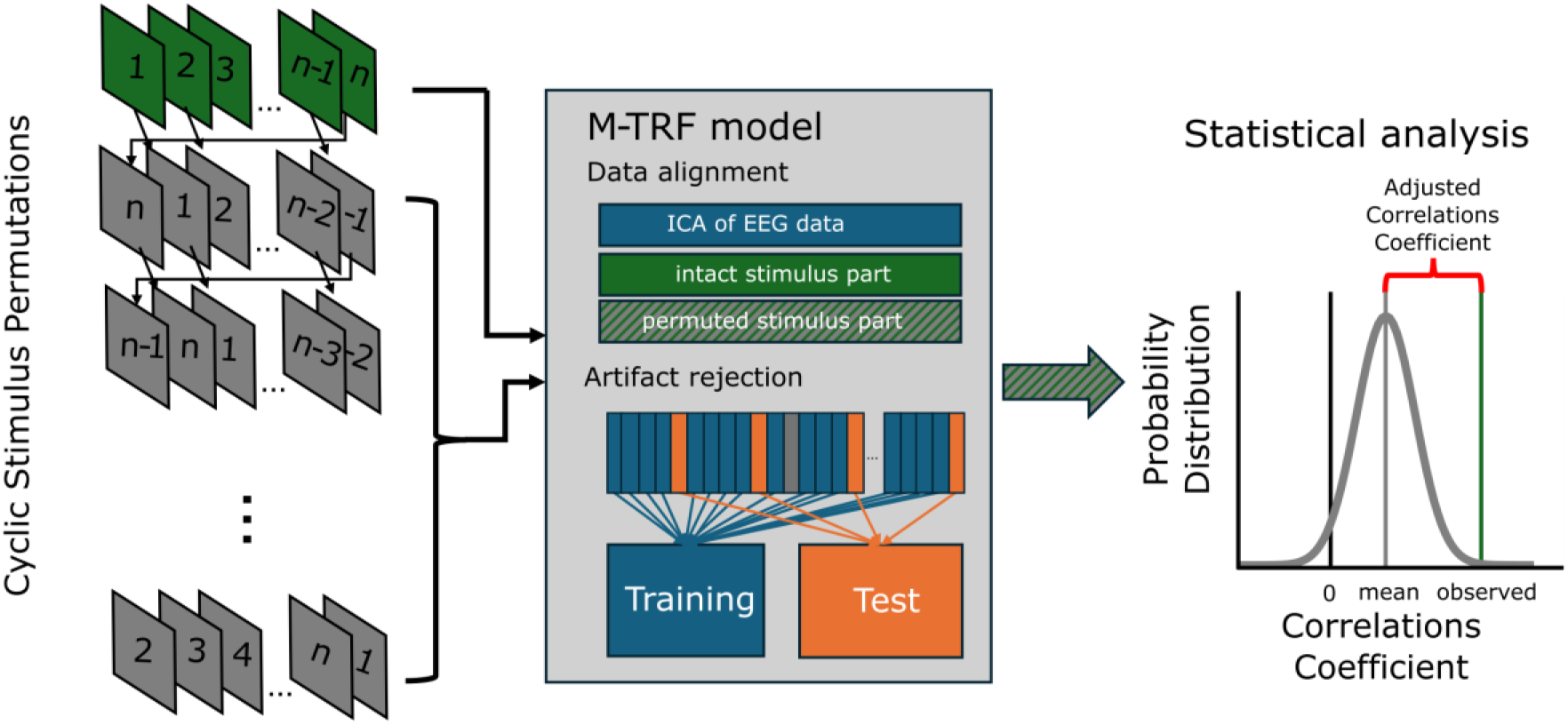
overview of the cyclic permutation analysis. On the left, a schematic illustrates how the stimulus sections are rearranged across the different permutations. The middle panel outlines stimulus alignment and the m-TRF model’s input partitioning. On the right, a schematic histogram shows the distribution of correlation coefficients for the model trained using permuted stimuli. The mean of these histograms quantifies the correlation that can be explained without the cyclically permuted variables. The red brace indicates the correlation exclusive to the permuted variables.

In this study, we applied 100 cyclic permutations, spaced equally in time. This results in a spacing between cycles of the permutations of ∼ 4 seconds. Notably, the permutation process was performed without regard to rejected epochs, which ensures perfect equality between the real data sets and the surrogate data sets. Selecting the number of cyclic permutations involves balancing two factors: obtaining enough surrogate-datasets for robust statistical analysis, while simultaneously ensuring that each permutation is spaced far enough apart in time from its neighbors to be considered statistically independent.

To each of these cyclic permutations, an mTRF model is fitted using the numerically estimated ridge parameter. For each model, the correlation coefficient between the model’s prediction and the data is calculated and averaged across all ICA components. The cyclic permutation analysis comes with four key benefits:

**1. quantifying overfitting**. The degree of overfitting is estimated as the mean correlation coefficient explained by the 99 (N-1) cyclic permutations that are not the correctly time-aligned stimulus. Compensating for this overfitting is as simple as calculating the stimulus-related correlation as the difference between the total correlation observed in the model with correctly time-aligned stimulus and the mean correlation observed using the surrogate data sets as inputs.
**2. additional objective data rejection criterion**. One concern with EEG recordings is that they are noise-sensitive measurements; therefore, there is significant variation in data quality, and some datasets cannot be successfully analyzed, regardless of the data preprocessing or modeling approach. These instances of poor data quality can be systematically identified using the variance of overfitting as a measure of data quality (meaning the variance of the cyclic permutation when all input variables are permuted). In the present study, all stimulus stories with an overfitting variance of more than five standard deviations outside of the mean of the variance of the accepted stories were excluded.

We also retroactively applied this rejection criterion to the conventional mTRF model to ensure that observed differences between the models are not just an artifact of the improved rejection.

**3. selective removal of information from the input**. This is done by permuting only a subset of input variables, then calculating the correlation attributed to them as described above. It should be noted that the mean correlation of the models based on the surrogate data set could now be significantly larger than zero and no longer measures overfitting, but rather the sum of overfitting and the correlation that can be explained by the part of the input space that has not been permuted.
**4. empirically derived p-value**. We directly measured the statistical certainty for additional explained variance, without the need to compare models of differing input sizes. This was done by first confirming the model’s correlation coefficients are normal, then using the mean and variance of the incorrect cyclic permutation to compute the z-transform of the observed signal, and finally converting the resulting z-values into p-values.

### Simulation of the lambda parameter

Conventionally, the optimal ridge parameter can be determined through cross-validation. As mentioned above, this comes at both a computational cost and introduces additional susceptibility to noise. Here we propose an alternative approach of estimating it through numerical simulation. Concretely, the ridge parameter is estimated by generating multiple surrogate-datasets as described below, and an mTRF model is fitted at each candidate lambda (34 logarithmically spaced values, ranging from 10^-2^ to 10^9^). Correlation coefficients for each lambda are averaged across all surrogate-datasets, and the one yielding the highest average correlation is selected as optimal.

The generation of surrogate data sets is done by generating each surrogate signal based on a different cyclic permutation of the stimulus to which noise is added. This ensures that the selection of lambda does not bias the mTRF toward any particular cyclic permutation and is particularly important when only a subset of the input variables is permuted. The surrogate signal is simply a convolution of the mTRF lag times matrix of the stimulus with a random matrix shaped like the kernel of the mTRF model. In total, 36 surrogate data sets and normally distributed noise with a root-mean-square amplitude 16 times that of the surrogate signal were analyzed.

Model fitting to each dataset was done using the same procedure as described in the previous section. The only difference is that we can use the noise-free surrogate signal in calculating the correlation coefficient, which dramatically reduces our susceptibility to the effects of noise.

### Evaluation of the reliability of the models

To evaluate whether the transition to the ICA space and the partitioning into shorter segments can enhance the model’s performance, one key metric is the reliability of cross-validation in determining the optimal lambda. This was assessed by examining the distribution of the location of the highest slope in the correlation coefficient as a function of the ridge parameter and comparing their distributions using a Kolmogorov-Smirnov test.

#### EEG Data collection

An active listening task was completed during the EEG measurement. Participants listened to six audio stories, each approximately six minutes long, followed immediately by six comprehension questions per story to ensure vigilance. All stories were narrated by the same male voice actor and delivered monophonically at a comfortable volume using a free-field loudspeaker. The six stories were designed to evoke different emotions, including a funny story, a fantasy story, a boring story, a realistic story, an adventure story, and a second boring story, in that order. For a full description, please see [26]. Transcripts and recordings of these materials are available at https://osf.io/hnc4f EEG recordings took place in a sound-attenuating, and electrically shielded booth. The participants were seated comfortably and instructed to minimize movement and look at a book icon on the screen in front of them, serving as a fixation cross. The audio stimuli were delivered at a sampling rate of 48,000 Hz using a Rokit 5 KRK loudspeaker located in front of the participant.

EEG data were collected with a 64-channel BioSemi ActiveTwo EEG system at 512 Hz. Before recording, the offset voltage for each electrode relative to the ground electrode was verified to be below 20 µV to ensure data quality and accuracy. To properly align the EEG with the auditory stimulus and ensure robustness against a potential clock drift, the second audio channel (not audible to participants) was used to send a trigger to the EEG system every minute. Linear regression of the observed trigger times against the expected trigger times revealed that the clock drift was minimal, at approximately 4 ms per minute, and thus falls within the specifications of the EEG system and sound card. Furthermore, the observed temporal accuracy exceeded the alignment required for the mTRF analyses, which were performed at sampling rates of 64 Hz.

#### EEG preprocessing

EEG data were preprocessed using a custom MATLAB script (version 2025a) using the EEGlab toolbox (v2025.1.0). First, a 4th-order bidirectional Butterworth bandpass filter was applied, retaining frequencies between 1 and 8 Hz, which results in a −24 dB/octave roll-off at the edge. Next, the data were downsampled to 64 Hz to enhance computational efficiency. Subsequently, detrending was performed by subtracting the mean calculated over a 5-second sliding window from each time point.

The next step was the automatic removal of artifacts. The process involved dividing the data into 1-second epochs. All epochs with an average variance across all channels that deviated from the mean by more than five standard deviations of the variances of the accepted epochs were removed as artifacts. Considering only the accepted epochs to estimate the mean variance ensures stability against outliers. In practice, the mean and standard deviation of variance were iteratively calculated (100 iterations).

Finally, independent component analysis (ICA) was performed using the ‘runica’ ICA algorithm to decompose the EEG signals into statistically independent components. The ICA was performed independently for each participant and story, and used only the included epochs.

#### Recruitment exclusion and inclusion criteria

The participants were recruited from the University participation pool and via word of mouth. All participants were compensated for their time (16 CAD per hour). All methods were reviewed by the University of Western Ontario Nonmedical Research Ethics Board (REB#125776). All participants provided written and informed consent before participating.

### Stimulus preprocessing

We considered two types of variables for modeling the input space. The first captures auditory information by decomposing the acoustic speech signal into a series of band-pass filters mimicking the human ear. The second type encodes phonetic information in which constituent phonemes are coded as binary vectors that denote linguistic features.

The filtering of the incoming acoustic waveform via the anatomy of the inner ear was artificially recreated using the Equivalent Rectangular Bandwidth (ERB) scale, which divides the auditory frequency range into perceptually equivalent bands [27]. In studies examining the correlation between EEG and speech spectra, it is common to use bands that are 1.6 ERB wide [11,28]. Following this convention, we employed 19 bands of 1.6 ERB width, spanning 0.25 to 8.2 kHz. This approach ensures coverage of the frequency range most critical for speech comprehension.[29]

For each frequency band, we extracted the amplitude envelope by taking the absolute value of the Hilbert transform. To prevent aliasing during downsampling to the EEG sampling rate, each envelope was subsequently filtered with a low-pass filter at the Nyquist frequency.

Phoneme onsets for the speech stimuli were extracted using the Montreal Forced Aligner [30] at an accuracy of ±10ms. Next, each phoneme was represented by binary features denoting individual phonetic features such as voicing, place, and manner of articulation for consonants, and height, rounding and tongue position for vowels (see [31] for details).

